# Transglobal spread of an ecologically significant sea urchin parasite

**DOI:** 10.1101/2023.11.08.566283

**Authors:** Isabella T. Ritchie, Brayan Vilanova-Cuevas, Ashley Altera, Kaileigh Cornfield, Ceri Evans, James S. Evans, Maria Hopson-Fernandes, Christina A. Kellogg, Elayne Looker, Oliver Taylor, Ian Hewson, Mya Breitbart

## Abstract

Mass mortality of the dominant coral reef herbivore *Diadema antillarum* in the Caribbean in the early 1980s led to a persistent phase shift from coral-to algal-dominated reefs. In 2022, a scuticociliate most closely related to *Philaster apodigitiformis* caused further mass mortality of *D. antillarum* across the Caribbean, leading to >95% mortality at affected sites. Mortality was also reported in the related species *Diadema setosum* in the Mediterranean in 2022, where urchins experienced gross signs compatible with scuticociliatosis. However, the causative agent of the Mediterranean outbreak has not yet been determined. In April 2023, mass mortality of *D. setosum* occurred along the Sultanate of Oman’s coastline. Urchins displayed signs compatible with scuticociliatosis including abnormal behavior, drooping and loss of spines, followed by tissue necrosis and death. Here we report the detection of an 18S rRNA gene sequence in abnormal urchins from Muscat, Oman that is identical to the *Philaster* strain responsible for *D. antillarum* mass mortality in the Caribbean. We also show that scuticociliatosis signs can be elicited in *D. setosum* by experimental challenge with the cultivated *Philaster* strain associated with Caribbean scuticociliatosis. These results demonstrate the *Philaster* sp. associated with *D. antillarum* mass mortality has rapidly spread to geographically distant coral reefs, compelling global-scale awareness and monitoring for this devastating condition through field surveys, microscopy, and molecular microbiological approaches, and prompting investigation of long-range transmission mechanisms.

## Introduction

The long-spined sea urchin genus *Diadema* is ubiquitous in tropical reef habitats across the globe, where it exerts critical control on algal growth (1), allowing sufficient light and space for new corals to settle and thrive (2, 3). The loss of these important herbivores can result in phase shifts from coral-to algal-dominated communities, with widespread ecosystem effects (1, 4).

A mass mortality event of unknown etiology decimated Caribbean *Diadema antillarum* populations in the early 1980s, with very limited recovery in subsequent years (4–7). Another *D. antillarum* mass mortality event was first reported in February 2022 in the U.S. Virgin Islands and by May 2022 abnormal urchins were observed across the Caribbean (7, 8). The 2022 mass mortality was caused by a scuticociliate most closely related to *Philaster apodigitiformis* (8). Signs of the condition, termed *Diadema antillarum* scuticociliatosis (DaSc), include abnormal behavior, loss of tube foot control, stellate spine orientation, spine drooping and loss, and finally tissue necrosis and death (7, 8). In both the 1980s and 2022 die-offs, no other echinoid species were affected (4, 8).

Beginning in July 2022, mass mortality was observed in clade b *Diadema setosum* in its invasive range in the Mediterranean Sea (9). Signs resembled DaSc (8, 9), but the etiological agent was not determined. Here we demonstrate the presence of *P. apodigitiformis* in abnormal urchins from Muscat, Oman, establishing the spread of scuticociliatosis to the native range of clade b *D. setosum* in the Sea of Oman, and show that *P. apodigitiformis* can elicit scuticociliatosis in clade a *D. setosum*.

## Results

In April 2023, we observed abnormal clade b *D. setosum* in the Sea of Oman (Figure 1). *P. apodigitiformis* was detected by nested PCR (10) in six abnormal urchins collected from the Capital Area Yacht Club in Muscat, Oman. The 18S rRNA gene sequences from these samples were identical to *P. apodigitiformis* FWC2 cultured from abnormal *D. antillarum* in the Caribbean (8) (Figure 2).

**Figure 1.**
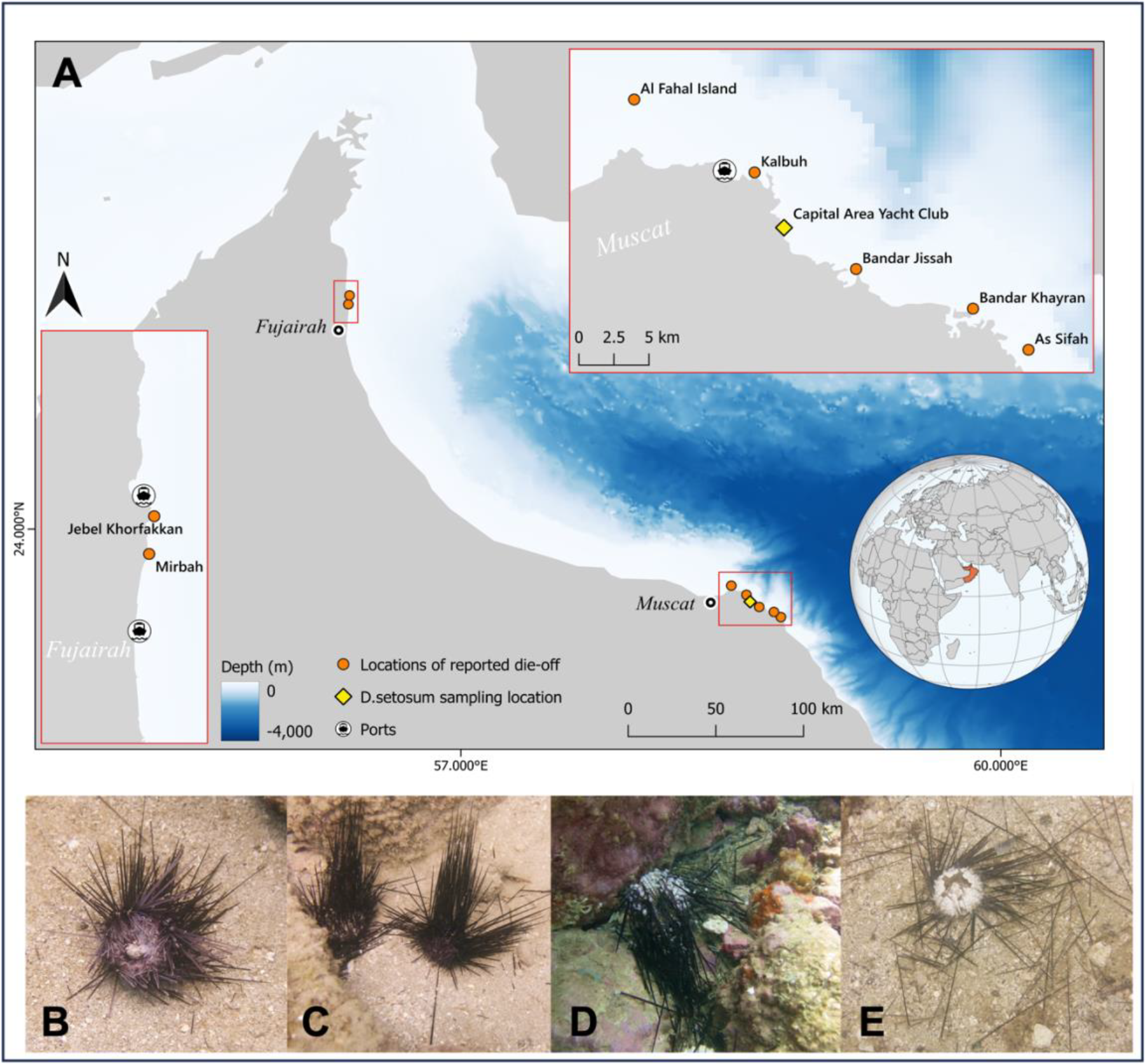
Abnormal *Diadema setosum* reported from the Sea of Oman. (A) Graphical depiction of locations in the Sea of Oman where *D. setosum* die-offs were reported (orange points) and confirmed in collected individuals via PCR (yellow diamond). Inlaid images show more detailed maps of sites in Fujairah (left) and Muscat (top). Basemap created in QGIS using data freely available from General Bathymetric Chart of the Ocean (www.gebco.net) and EOX (maps.eox.at). (B-E) Images depicting signs seen in urchins considered ‘grossly abnormal’, including unusual behavior (B), stellate spine arrangement (C), spine loss and tissue necrosis (D), and eventual death (E). Photo credit: Kaileigh Cornfield.

**Figure 2.**
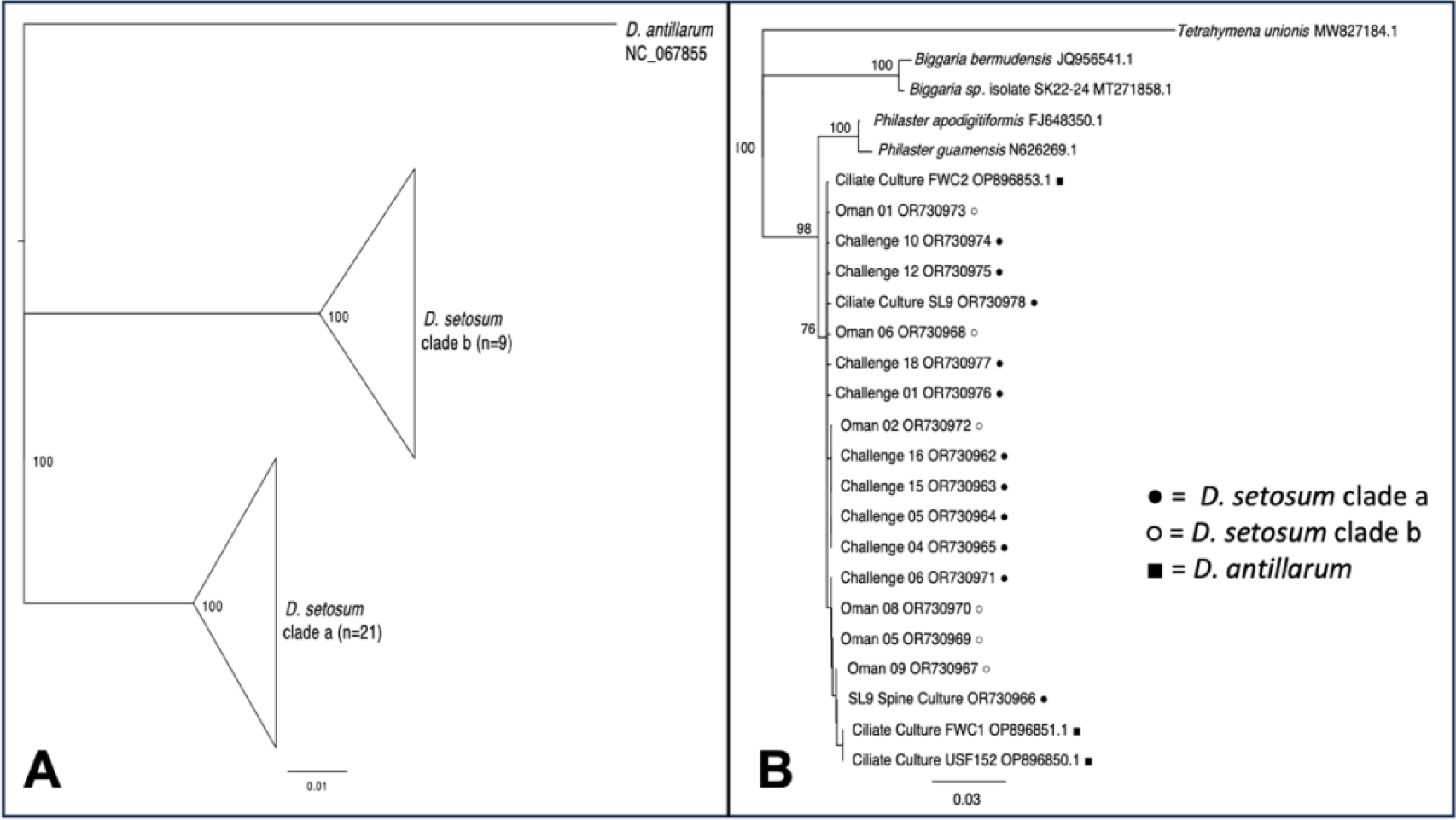
*Diadema* and scuticociliate phylogenies. (A) Phylogenetic representation of *D. setosum* clades a and b sampled during this experiment, as well as *D. antillarum*. (B) Phylogenetic representation of scuticociliate sequences from *D. setosum* (clade a = filled circle, clade b = empty circle), *D. antillarum* (closed square), and close relatives identified by BLASTn searches.

To determine if clade a *D. setosum* is also susceptible to scuticociliatosis, we ordered specimens from commercial aquarium suppliers for use in challenge experiments. One urchin (SL9) presented scuticociliatosis signs upon arrival and we observed ciliates resembling *P. apodigitiformis* swarming in dropped spines under microscopy, leading us to establish a culture from this urchin’s coelomic fluid. PCR and sequencing confirmed 100% identity between the 18S rRNA gene sequences of the FWC2 and SL9 cultures (Figure 2). Detecting this scuticociliate in an urchin obtained through the aquarium trade provided initial evidence for the ability of *P. apodigitiformis* to infect clade a *D. setosum*, leading us to conduct a controlled five-day experimental challenge.

Following established protocols (8), six urchins were challenged with FWC2, six with SL9, and six served as negative controls. Five of the six urchins treated with each ciliate culture lost spines and died, while only two of the six controls (one each of FWC2 and SL9 filtrate) exhibited signs of infection, likely resulting from ciliate exposure prior to arrival at our facility (Figure 3A). Grossly normal urchins had lower levels of *P. apodigitiformis* in body wall, spine, and coelomic fluid samples than abnormal urchins by quantitative PCR (qPCR) for the 28S rRNA gene, regardless of treatment (Figure 3B).

**Figure 3.**
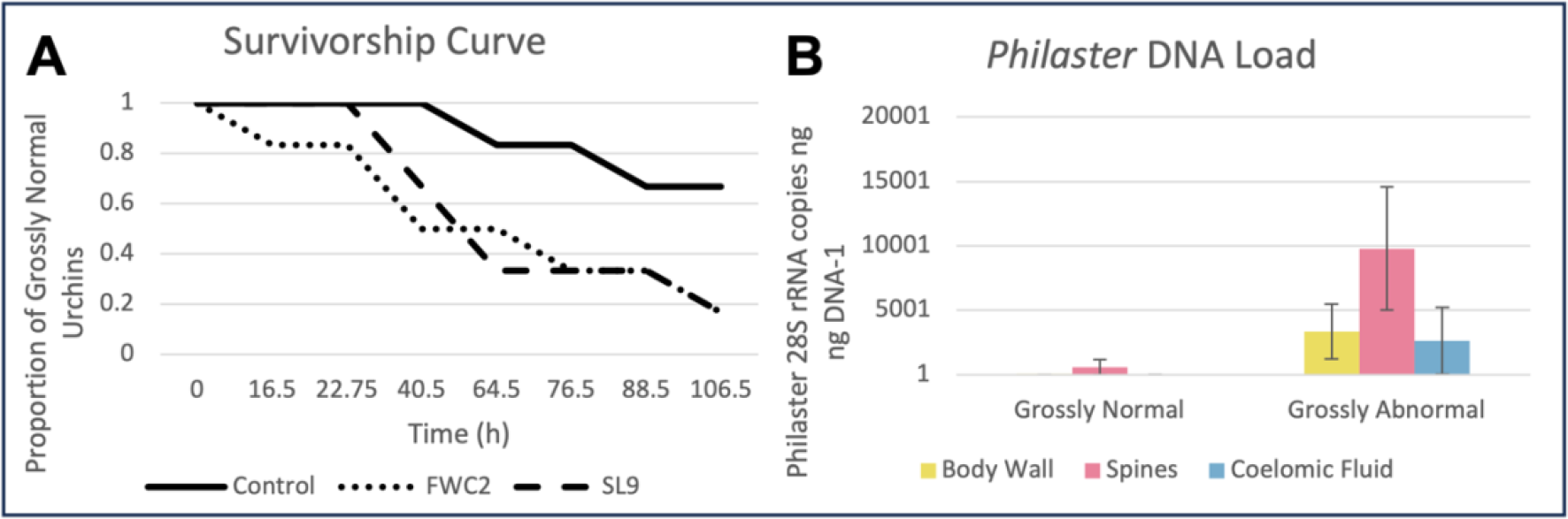
Survivorship curves and *Philaster* load for experimentally challenged clade a *Diadema setosum*. (A) Survivorship curve showing the decrease in grossly normal individuals over time for urchins treated with two *Philaster* cultures (FWC2 and SL9) and the controls. (B) Quantitative PCR results for body wall, spines, and coelomic fluid samples from the 18 experimental challenge urchins, classified by animal state at the end of the experiment.

## Discussion

Our experimental challenge results combined with field samples from Oman demonstrate that *P. apodigitiformis* can cause scuticociliatosis in both clades of *D. setosum*, representing a significant threat to these important herbivores. While *D. setosum* is an invasive species in the Mediterranean Sea, the detection of scuticociliatosis in the Sea of Oman, part of its native Red Sea habitat, could have disastrous ecological effects on coral reef communities reminiscent of those seen in the Caribbean following the 1980s die-off (1, 4, 6).

These results highlight the importance of monitoring urchin densities and ecosystem-level effects of loss of these keystone herbivores in affected regions. Additionally, our experimental infection of clade a *D. setosum* indicates the vulnerability of this population to scuticociliatosis should the ciliate reach its native range, emphasizing the need for baseline benthic surveys. We also observed mortality of *Echinothrix* sp. in Fujairah and Al Fahal Island, suggesting that other species in the *Diadematidae* family are susceptible to scuticociliatosis.

While the negative impacts of scuticociliatosis on *Diadema* spp. are clear, many questions remain about the factors that affect *P. apodigitiformis* growth and pathogenicity and *Diadema* immune responses. Although similar signs of scuticociliatosis were observed in *D. antillarum* and *D. setosum*, individuals of both species display variability in their responses to infection (including some experimentally infected specimens of both species that remained grossly normal). Finally, now that *P. apodigitiformis* has been detected in geographically disparate locations, it is critical to assess long-range transmission routes of this ciliate, including natural (e.g., currents, seabirds) and anthropogenic (e.g., shipping vessels, recreational diving, aquarium trade) pathways.

## Materials and Methods

DNA was extracted from urchin samples (∼1 mm fragments of the body wall, 1-3 spines with bases, or 200 μl of coelomic fluid) using the Zymo Quick-DNA Tissue/Insect Kit following manufacturer’s instructions with the exceptions of omitting β-mercaptoethanol from the lysis buffer, bead-beating for 2 minutes, and eluting into sterile water. Nested PCR was used to specifically amplify and sequence the *Philaster* clade associated with *Diadema* scuticociliatosis (10, 11). Urchin species and clade identification was confirmed through CO2b/ATP6b amplification and sequencing (12). All ciliate sequences generated in this study are available in GenBank (Accession Numbers: OR730962-OR730978). Phylogenetic analyses were performed in Geneious Prime using the Geneious Tree Builder and Tamura-Nei genetic distance model with the neighbor-joining method.

FWC2 was cultured from the coelomic fluid of an abnormal *D. antillarum* collected in the Florida Keys in June 2022 and has been maintained in xenic culture. SL9 was cultured from an abnormal clade a *D. setosum* obtained through the aquarium trade by drawing 0.5 ml of coelomic fluid into a sterile syringe fitted with a 22-gauge needle by insertion of the needle through the peristomal membrane and inoculating into 5 ml of media composed of 0.2 μm-filtered seawater, 0.01% yeast extract, an autoclaved grain of rice, and 250 μl 0.2 μm-filtered *D. antillarum* extract (8). Within 48 h of incubation at room temperature, the SL9 culture was densely populated by ciliates similar in morphology to FWC2 and dilution-to-extinction was performed.

For the challenge experiment, eighteen urchins acquired through aquarium suppliers were placed into individual tanks filled with ∼7 liters of 5 μm-filtered offshore Florida Keys seawater and an airstone bubbler. Twelve urchins were inoculated with ∼250 ciliates each by addition to the water directly above the urchin (n=6 treated with FWC2, n=6 treated with SL9), and the remaining six urchins were treated with the same volume of 5 μm-filtered culture (n=3 FWC2, n=3 SL9) to control for bacteria within the media. Urchins were monitored for signs of infection and collected when disease was apparent or at experiment termination after five days. Upon collection, urchins were dissected to obtain coelomic fluid, spine/spine base, and body wall samples, which were frozen at -80°C until DNA extraction and quantitative PCR for *P. apodigitiformis* following previously published protocols (8).

## Acknowledgments

This work was supported by the Atkinson Center for Sustainable Futures Rapid Response Fund and OCE-2049225 (IH) and the Von Rosenstiel Fellowship and Von Rosenstiel Innovation Fund for Marine Science (ITR). ITR and BVC were supported by the NSF Graduate Research Fellowship Program (#2136515 and #1650441). JSE and CAK were supported by the U.S. Geological Survey Ecosystems Mission Area Biological Threats and Invasive Species Research Program.

Any use of trade, firm, or product names is for descriptive purposes only and does not imply endorsement by the U.S. Government.

